# Bioengineering of genetically encoded gene promoter repressed by flavonoids for constructing intracellular sensor for molecular events

**DOI:** 10.1101/2021.02.17.431641

**Authors:** Nicole M. Desmet, Kalyani Dhusia, Wenjie Qi, Andrea I. Doseff, Sudin Bhattacharya, Assaf A. Gilad

## Abstract

In recent years, Synthetic Biology has emerged as a new discipline where functions that were traditionally performed by electronic devices are replaced by “cellular devices”. Those are genetically encoded circuits, constructed of DNA that are built from biological parts (aka bio-parts). The cellular devices can be used for sensing and responding to natural and artificial signals. However, a major challenge in the field is that the crosstalk between many cellular signaling pathways use the same signaling endogenous molecules that can result in undesired activation. To overcome this problem, we utilized a specific promoter that can activate genes with a natural, non-toxic ligand at a highly-induced transcription level with low background or undesirable off-target expression. Here we used the orphan aryl hydrocarbon receptor (AHR), a ligand-activated transcription factor that upon activation binds to specific AHR response elements (AHRE) of the Cytochrome P450, family 1, subfamily A, polypeptide 1 (CYP1A1) promoter. Flavonoids have been identified as AHR ligands. Data presented here shows successful creation of a synthetic gene “off” switch that can be monitored directly using an optical reporter gene. This is the first step towards bioengineering of a synthetic, nanoscale bio-part for constructing a sensor for molecular events.

## Introduction

Recent innovations in Synthetic Biology, including dramatic reductions in the cost of DNA synthesis and sequencing [1,2], opened many new possibilities for using biological material as a replacement for traditional “electronic devices”. As such, biological components can be combined with electronic biosensors for miniaturing, and to improve integration with target tissue. For such integration to occur, a new set of biological parts (or “*bio-parts*”) should be developed. Those bio-parts are required to be highly specific and have the ability to communicate with electronics. The input and output is commonly achieved by light [3–7], chemicals [8,9] or electromagnetic irradiation [10–15].

The idea that genes could be activated or suppressed was first pioneered in prokaryotic cells due to their ease and simplicity, but gene switches have since been developed for eukaryotic cells as well. Gene switches are natural or synthetic systems that allow initiation, interruption or termination of target gene expression [16]. Many of the gene switches are based on gene promoters. Promoters consist of DNA sequences usually preceding the gene open reading frame which are essential for gene regulation. There are only few examples that are standing out for the use of promoters in mammalian cells as switches, that are activated by metabolites[17], ions[18], optogenetics[19] or miRNA[20], but essentially the most effective gene switch is tetracycline-dependent repressor (TetR)[21]. This promoter/repressor system is the most ubiquitous system that is used in mammalian cells. Mainly because it has no crosstalk with other signaling pathways, it does not activate any other gene and consequently, has no cellular side effect. In this study, we sought to expand the synthetic promotor toolbox available as bio-parts for future design and construction of bio-electronic hybrid devices.

The aryl hydrocarbon receptor (AHR) is a ligand-activated transcription factor that serves as a sensor of developmental and environmental signals [22] but whose endogenous activators and their roles are poorly understood [23]. Inactivated, the AHR resides in the cytosol bound to inhibitory cofactors, including heat shock protein 90 (hsp90) and aryl hydrocarbon receptor interacting protein (AIP) (Figure 1). Upon activation by a ligand, the cytosolic AHR undergoes a conformational change that frees it from the inhibitory cofactors [24]. This allows the AHR-ligand complex to translocate into the nucleus and associate with the aryl hydrocarbon receptor nucleus translocator (ARNT) protein [25]. This complex then binds to specific DNA sequences on target genes called AHR response elements (AHRE) and is known to activate the transcription of the Cytochrome P450, family 1, subfamily A, polypeptide 1 (CYP1A1) by binding specifically to its upstream promoter.

**Figure 1.**
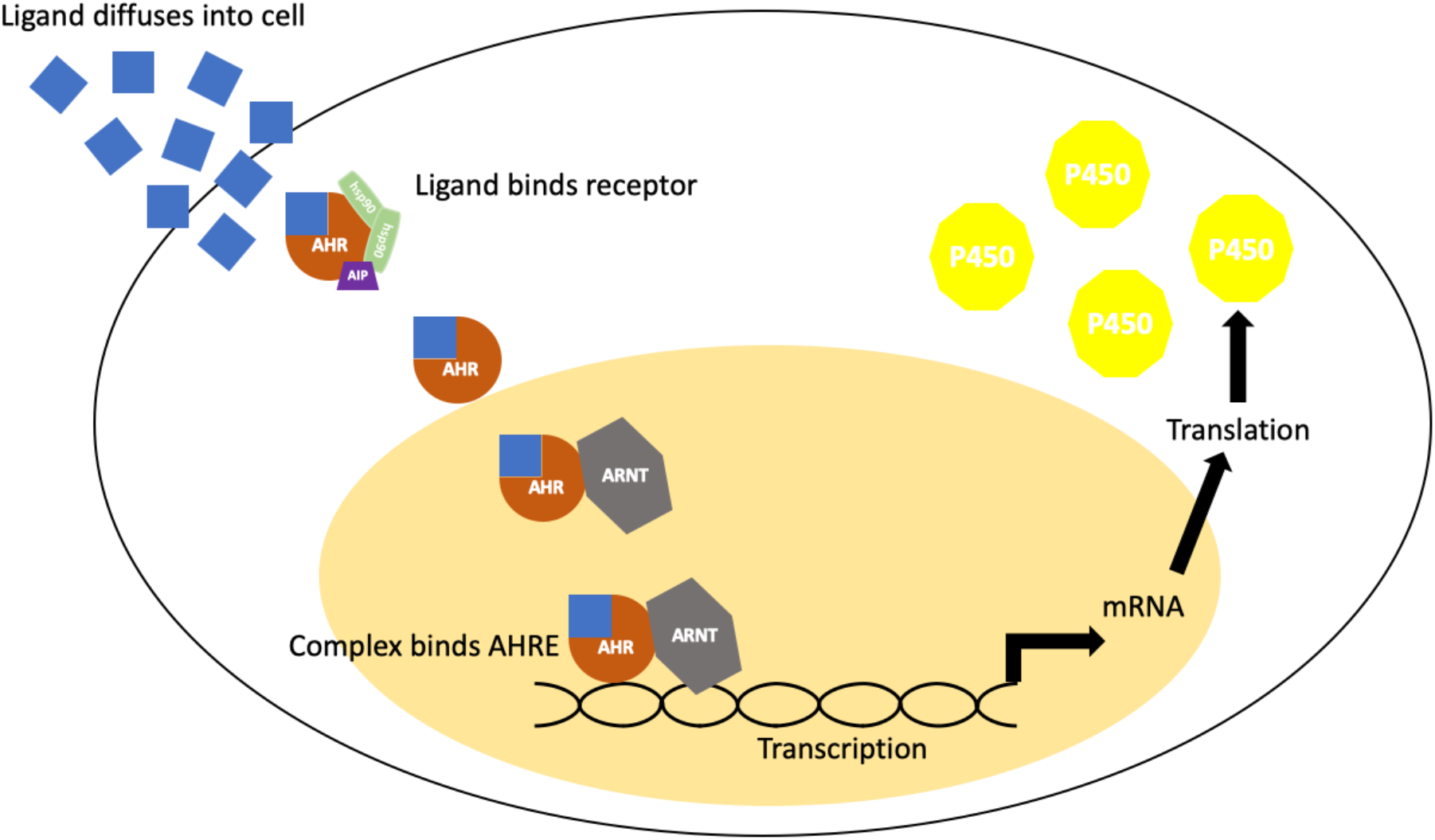
AHR/CYP1A1 pathway diagram.

Most known AHR ligands are toxins and xenobiotics, such as 2,3,7,8-tetrachlorodibenzo-B-dioxin (TCDD) [24]. Additionally, flavonoids, one of the most abundant phytochemicals, have been identified as weak AHR ligands [26]. Flavonoids are a class of polyphenolic secondary plant metabolites broadly found in fruits and vegetables. These compounds are generally recognized as health promoting, immuno-modulators and are components of traditional medicines and commercially-available nutraceuticals [27].

This gene circuit, while complex, gives three accessible points of modification: the ligand, the receptor, and the promoter. In this study, we discussed our work on each, and how these come together to produce a sensitive synthetic gene switch which can be remotely suppressed, as well as future work for developing a sensitive activator. The applications of this cellular system are broad and include immune or stem cell imaging and tracking, *in situ* remote activation of gene expression in cells of interest as well as implementing Synthetic Biology in building bio-electronic hybrid sensors.

## Materials and Methods

### Plasmid synthesis and construction

A 2 Kb region upstream of the *CYP1A1* gene is in the human genome part of the promoter region. This sequence was used as the experimental CYP1A1 promoter. To determine an optimal promoter sequence that will drive the expression of a reporter gene in response to flavonoids, a machine learning model, XGBoost [28], was trained by the AHR binding sites found in human breast cancer MCF-7 cells. The promoter region was cloned into pGlow TOPO TA expression plasmid (Invitrogen, catalog number: K483001) following the product’s protocol. Additionally, the reporter gene GFP was replaced with mScarlet using the NEBuilder HiFi DNA Assembly kit (New England BioLabs catalog number:E5510S), to produce two different expression vectors.

### Cell Culture

HeLa and HEK293FT cells were cultured in Dulbecco’s modified Eagle’s medium (DMEM) (GIBCO catalog number:11965118) supplemented with 10% (v/v) fetal bovine serum (GIBCO catalog number 16000044) and 1% penicillin/streptomycin (GIBCO catalog number:15140122) at 37 °C in a humidified incubator with 5% CO_2_.

### Transfection

Cells were seeded in a 24-well plate at a density of 10^5^ cells/well and given overnight to properly adhere and subsequently transfected with either the experimental plasmid (pGlow-CYP1A1_prom_ or pGlow-CYP1A1_prom_::mScarlet), negative control plasmid (pcDNA3.1/V5-His-TOPO/lacZ), or no DNA using Lipofectamine 3000 Transfection Reagent (Invitrogen catalog number: L3000015) following the product protocol. Cells were incubated with the transfection mixture for 24 hours.

### Flavonoid Treatment

Transfected cells were then seeded in a 96-well plate at a density of 15,000 cells/well and given overnight to properly attach to the well bottom. Quercetin was purchased from Cayman Chemical (catalog number: 10005169) and dissolved in DMSO to a 220 mM stock solution (Sigma Aldrich catalog number: D8885-500G), apigenin and naringenin were purchased from Sigma and dissolved in DMSO as 20 mM and 100 mM stock solutions, respectively. Flavonoid stock solutions were kept at −20°C. Flavonoids were diluted in cell media to achieve 100 μ M working concentration, and then each flavonoid, in addition to a 0 μM control (DMSO), was added to each transfected cell group for 48 hours. This produced 12 individual experimental groups.

### Imaging

Following incubation, cells were washed with PBS (GIBCO catalog number: 10010023) and covered with Fluorobrite DMEM (GIBCO catalog number: A1896702). Images were taken on the Keyence BZ-X800 microscope using the BZ-X GFP filter for the GFP construct or the BZ-X TRITC filter for the mScarlet construct.

### Quantitative analysis

Following imaging, quantitative analysis was done using the VictorNIVO (PerkinElmer) microplate reader. The settings of excitation: 560 nm, emission: 593 nm were used.

### Structure prediction, docking and validation

The amino acid sequence of target protein 4xt2 named as crystal structure of the high affinity ARNT C-terminal PAS domains in complex with a tetrazole-containing antagonist was taken from the Protein Data Bank database (https://www.rcsb.org/structure/4XT2) [29]. To generate a reliable 3d structure 4xt2_A was remodeled using i-tessar target-template alignments.

To improve the quality of predicted model of 4xt2_A, energy minimization was performed with the gromos 96 force field. This force field permits to evaluate the energy of the modelled structure as well as overhaul distorted geometries through energy minimization. All computations during energy minimization were done in vacuum system, without relative force fields.

To assess flavonoid binding to the AHR (PDB: 4xt2), binding sites were predicted in the 3D structure of AHR using the CASTp server [30]. The interaction energy between the AHR and the ligand was scored for evaluation using Autodock Vina Package in mgltools. Line diagrams of flavonoids in AHR binding site generated by LIGPLOT [31]. The diagrams depict the hydrogen-bond interaction patterns and hydrophobic contacts between the ligand(s) and the main-chain or side-chain elements of the protein.

## Results and Discussion

A combined approach of molecular and computational biology was used to model and engineer a synthetic gene switch optimized in three ways: ligand, receptor, and promoter.

### Identifying the optimal promoter sequence

To determine an optimal promoter sequence that will drive the expression of a reporter gene in response to flavonoids, a machine learning model, XGBoost [28]. was trained on the AHR binding sites found in human breast cancer MCF-7 cells [32]. The resulting binding probabilities from 10 base pair segments containing the ‘GCGTG’ motif are displayed in Table 1. Screening of a 2Kb promoter region upstream of the transcription start site revealed 5 locations with putative binding probability of the AHR/ARNT complex greater than 0.5 suggesting that the complex bound to these locations would initiate transcription. This finding was validated by a computational model which shows AHR bound with ARNT form the transcription factor complex, visualized by UCSF Chimera Viewer (Figure S1). Therefore, the 2Kb sequence of the promoter was determined as sufficient and was used for cloning.

**Table 1.**
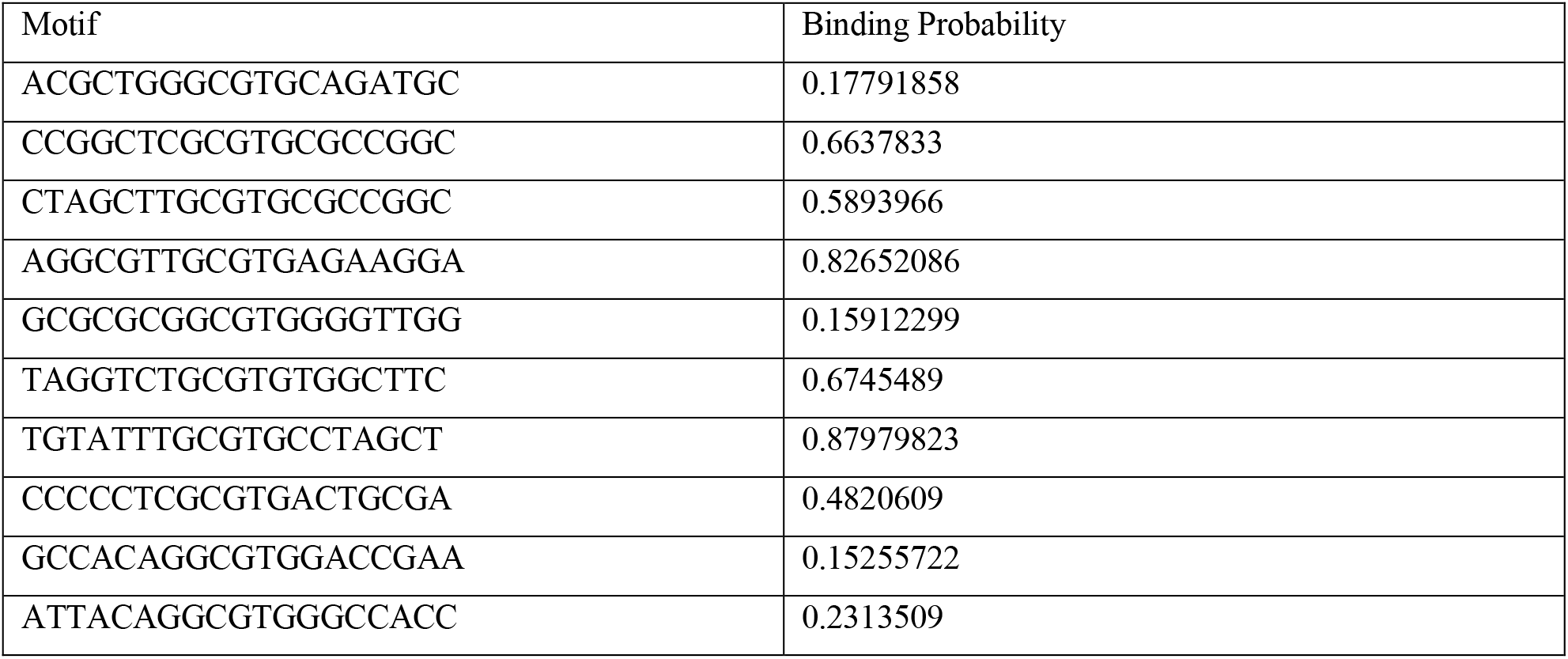
Ten putative AHR/ARNT binding sites and their corresponding binding probabilities.

### Bioengineering an optical reporter system

To determine the compatibility of the gene switch with the host cells, we showed that the AHR protein is expressed in HeLa and HEK293FT cells as confirmed by western blot analyses (Figure 2a). This step was crucial to ensure that these cells could support this reporter system. Next, a synthetic DNA fragment encoding to the 2Kb region of the human CYP1A1 promoter was cloned into the pGlow plasmid upstream of the EGFP reporter gene. Since flavonoids show autofluorescence in the green spectrum, EGFP was then replaced with a red shifted reporter, mScarlet, via Gibson assembly. The plasmid map is depicted in Figure 2b.

**Figure 2.**
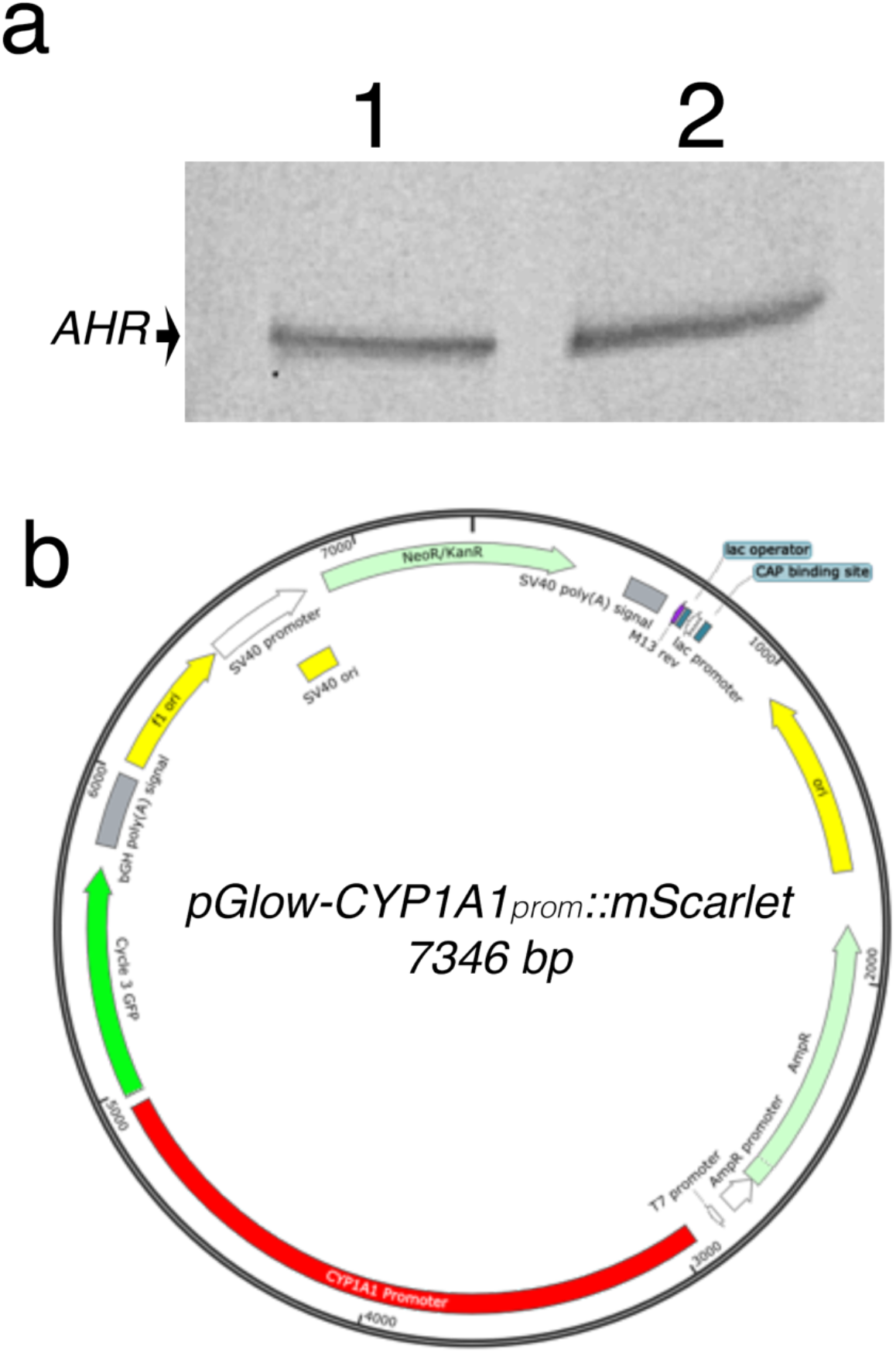
(**a**) Western blot analysis of AHR protein expression in HEK293FT (1) and HeLa (2). (**b**) Vector map of pGlow-CYP1A1_prom_::mScarlet plasmid.

### Testing the gene switch reporter system in vivo

To test the gene switch reporter system *in vitro*, HeLa cells were chosen for their stronger adherence to the plate, which was important due to the need of rigorous washes. After optimization of the timeline parameters (Supplementary material), cells were transfected with the pGlow-CYP1A1_prom_::mScarlet plasmid for 24 hours, then subsequently treated with 100 μM of either quercetin, apigenin or naringenin for 48 hrs. Interestingly, the pGlow-CYP1A1_prom_::mScarlet exhibits a basal level activation, which is eliminated by apigenin (Student’s t-test; p<0.01). Additionally, the signal was slightly enhanced by naringenin, and unchanged by quercetin, however, this was not found to be statistically significant. The control groups transfected with either pcDNA3.1/V5-His-TOPO/lacZ or No DNA exhibit no mScarlet expression, as shown in Figure 3. These findings indicate that among the three flavonoids tested, apigenin can be used successfully as a synthetic “off”-switch of gene expression.

**Figure 3.**
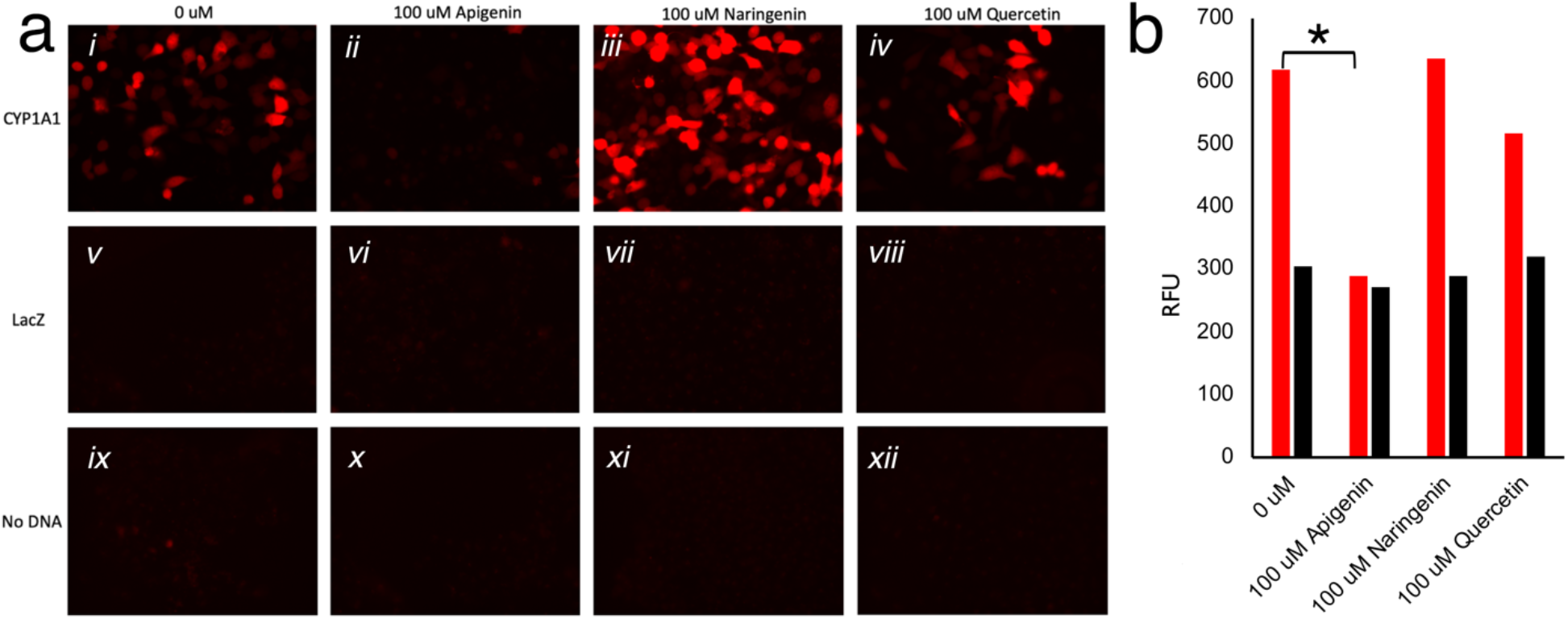
(***a***) HeLa cells imaged with TRITC filter. Twenty four hours post-transfection with CYP1A1_prom_::mScarlet, LacZ, or no DNA, cells were treated with 0uM (*i, v, ix* respectively), 100 μM apigenin (*ii, vi, x* respectively), 100 μM naringenin (*iii, vii, xi* respectively), or 100 μM quercetin (*iv, viii, xii* respectively) for twenty-four hours, washed, and imaged. (***b***) Fluorescence (Ex 560; Em 593) measured in HeLa cells transfected with CYP1A1_prom_::mScarlet or No DNA, treated with 0 µM, 100 µM apigenin, 100 µM naringenin, or 100µM quercetin. Red bars show significant reduction in the average fluorescence in the presence of apigenin, but not in other conditions, to the fluorescent level of untrasfected cells (black bars; * represents *p* < 0.01).

### Ligand/receptor binding analysis

To assess flavonoid binding to the AHR (PDB: 4xt2), binding sites were predicted in the 3D structure of AHR using the CASTp server, as shown in Figure 4.

**Figure 4.**
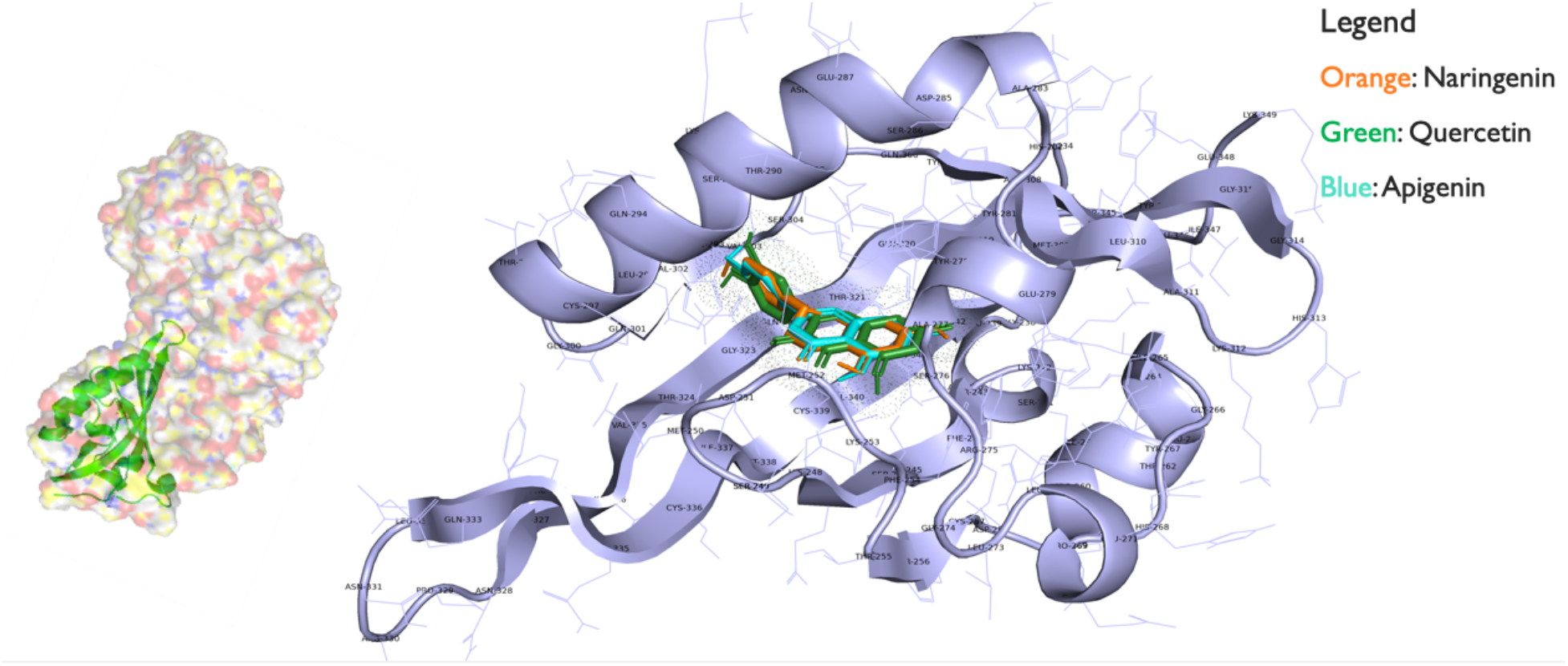
AHR bound with the flavonoids quercetin, naringenin and apigenin ligands using autodock from mgltools.

The interaction energy between the AHR and the ligand was scored for evaluation using Autodock Vina Package in mgltools. Results are shown in Table 2. The 43L or racemic tetrazolo-tetrahydropyrimidines are a natural substrate that bind with lower energy score, revealing that they have higher binding affinity towards 4xt2 (AHR) than TCDD. Ligand TCDD had the lowest energy score (−5.7 Kcal/mol) after 43L and apigenin which signifies that it has highest binding affinity towards the active site of 4xt2. The second and third best docked molecules were naringenin and quercetin which showed high binding affinity for the 4xt2.

**Table 2.** AHR ligands and their docking energy with AHR.

| Ligands | Binding energies (Kcal/mol) |
| --- | --- |
| Apigenin 5280443 <sup>1</sup> | -5.4 |
| Naringenin 932 <sup>1</sup> | -5.6 |
| Quercetin 5280343 <sup>1</sup> | -5.5 |
| 43L <sup>2</sup> | -5.8 |
| TCDD | -5.7 |

<sup>1</sup> PubChem number
<sup>2</sup> Inbound with protein

The results described above revealed the ability to generate a synthetic gene switch using the CYP1A1/AHR pathway. This pathway was chosen because it is dormant in mature cells and is not activated by any known endogenous ligand, only by exogenous –natural or synthetic-compounds. Flavonoids were screened to identify strong activators and suppressors of this gene circuit to allow for sensitive control. Thus far, apigenin was found to be an efficient suppressor. Using computational analysis, we found potential sites on the promoter region of AHR for future improvements of this gene switch. Promoter mutants were simulated and screened for the optimal sequence which would maximize transcription factor-promoter binding. Ligand-receptor binding simulations were done to identify key amino acids involved with binding in the ligand binding pocket of the AHR. These could be targets for protein mutagenesis to increase binding efficiency. Once all three prongs have been optimized, this fine-tuned synthetic gene switch could be utilized for vast imaging and therapeutic purposes.

While it is clear that we developed a reliable “off” system with apigenin, future optimization of the “on” system with naringenin is warranted. This would allow to increase the sensitivity without compromising on the tightness of the “off” switch. This could be accomplished by adding elements such as enhancers, co-factors, and genetic mutations to increase ligand-receptor binding strength and duration. Another direction to explore is to develop the “on” system by optimizing for a different flavonoid, such as luteolin or quercetin, and reducing the basal level.

Our overall goal is to engineer a completely synthetic signal transduction system that will allow controlling gene expression specifically and at will, while eliminating any crosstalk with other cellular components. This is essential for developing the future generation of *in vivo* Synthetic Biology – electronic devices hybrid biosensors.

## Acknowledgements

This project is supported in part by the NIH grants (for A.A.G) R01 NS098231, R01 NS104306 and P41 EB024495. We are thankful to Xiaxion Zhang for assistance with cloning and to Rita Martin for manuscript edits.

