## Supplementary material for "Bioengineering of genetically encoded gene promoter repressed by flavonoids for constructing intracellular sensor for molecular events": Supplamentary Information

HeLa cells were chosen versus HEK293FT cells because HeLa cells have stronger adhesive properties and therefore better handled the wash steps. To enhance efficiency, each trial was done in a 96-well microplate. Transfection protocol optimization consisted of varying cell concentration, transfection reagent concentrations, DNA concentration, and ratio to best enhance transfection efficiency. To determine the most efficient conditions, cells were transfected with pcDNA3.1/V5-His-TOPO/lacZ since the CYP1A1 plasmid is of similar size, and LacZ levels were quantified using a beta-galactosidase expression kit and microplate reader. Next, treatment conditions were optimized. In a time-lapse experiment, 48 hours was found to be the optimal treatment duration.

To simulate the ligand-receptor binding, the AHR/ARNT complex was used, shown in figure S1.

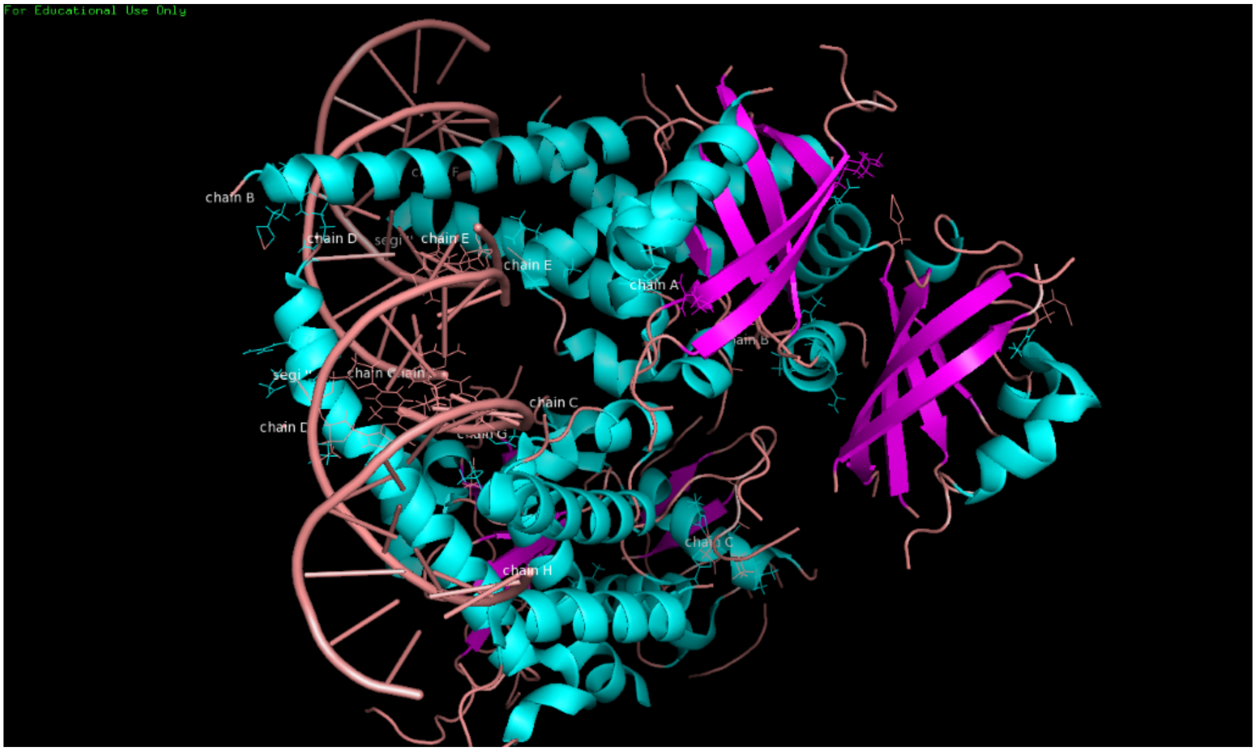


**Figure S1**. AHR/ARNT complex modeled by UCSF Chimera Viewer.
